# Fluctuation of cellular differentiation in limb regeneration is regulated by Pde4b in urodele amphibians

**DOI:** 10.1101/2021.09.13.460030

**Authors:** Akira Satoh, Rena Kashimoto, Ayaka Ohashi, Furukawa Saya, Sakiya Yamamoto, Takeshi Inoue, Toshinori Hayashi, Kiyokazu Agata

## Abstract

Urodele amphibians, *Pleurodeles waltl* and *Ambystoma mexicanum*, have organ-level regeneration capability, such as limb regeneration. Multipotent cells are induced by an endogenous mechanism in amphibian limb regeneration. It is well known that dermal fibroblasts receive regenerative signals and turn into multipotent cells, called blastema cells. However, the induction mechanism of the blastema cells from matured dermal cells was unknown. We previously found that BMP2, FGF2, and FGF8 (B2FF) could play sufficient roles in blastema induction in urodele amphibians. Here, we show that B2FF treatment can induce dermis-derived cells that can participate in multiple cell lineage in limb regeneration. We first established a newt dermis-derived cell line and confirmed that B2FF treatment on the newt cells provided plasticity in cellular differentiation in limb regeneration. Interspecies comparative analysis clarified that *Pde4b* upregulation by B2FF specifically took place in the newt cells. Blocking PDE4B signaling by Rolipram suppressed dermis-to-cartilage transformation and the mosaic knockout animals showed consistent results. Our results are a valuable insight into how dermal fibroblasts acquire multipotency during the early phase of limb regeneration *via* an endogenous program in amphibian limb regeneration.

---

Urodele amphibians can regenerate various organs including limbs. In their appendage regeneration, a structure called a blastema is induced after organ damage, and the induction of blastema is responsible for successful appendage regeneration. In other words, no blastema formation essentially results in no appendage regeneration. Thus, understandings of blastema formation would be the closest way to elucidate the ability of organ-level regeneration in urodele amphibians. And limb regeneration is an ideal model to study blastema formation and the organ-level regeneration ability in urodele amphibians.

Limb regeneration necessarily accompanies blastema formation, which has been considered to be a similar structure to a developing limb bud ^1^. A regeneration blastema actually showed a similar gene profile to a developing limb bud ^2^. Thus, a mechanism to induce a re-developmental field in an adult body has been investigated with great curiosity. From this angle, two issues are embossed: 1) how is the re-developmental field induced? 2) how do the differentiated cells turn into cells with an embryonic gene profile? Regarding blastema induction, classic studies clearly demonstrated that the nerves play a central role in the process ^3, 4^ Denervation from limbs results in no regeneration after limb amputation. Wound healing takes place instead. The amputation plane is covered by the migrating epidermis immediately after limb amputation. After the epidermal covering of the exposed amputation surface, the nerves secret some factors to create a regenerative environment. Molecules, which are secreted from the nerve ends and contribute to creating the regenerative environment in an amputated limb, had been investigated for a long time. Some factors have been identified as nerve factors ^5, 6, 7^. Among them, FGFs and BMPs can induce a blastema in multiple species and organs ^8^. *Fgf2*, *Fgf8*, and *Bmp2* are expressed axolotl neurons in the dorsal root ganglion (DRG). Application of FGF2+FGF8+BMP2 (B2FF) to the wounded skin results in blastema formation instead of skin wound healing in urodeles ^8^. FGF2+FGF8 without BMP2 (FF) can induce a blastema. However, the FF-induced blastema does not have the ability to keep growing up to a patterned limb. BMP2 induced structure lacks proper blastema gene expressions and does not have the ability to grow a limb ^8^. Downregulation of those genes in DRG neurons resulted in decreasing in the limb regeneration ability ^9^. Thus, FGF2, FGF8, and BMP2 can work as nerve-secreting molecules and regeneration inducers in urodele amphibians. Thanks to the determination of the inductive molecules, an approach to the other issue of how the differentiated cells turn into an embryonic state can be possible.

A regeneration blastema fulfilled by undifferentiated cells is induced on the amputation plane, below which differentiated tissues exist. Undifferentiated blastema cells, having an embryonic profile, emerged from the differentiated tissues ^2^. The dermis is the major source of blastema cells among the limb tissues ^10, 11^. Dermal fibroblasts have been considered to turn into blastema cells. Dermis-derived blastema cells can change their cell type into varied connective tissue lineage, such as cartilage. Such transdifferentiation ability is restricted within the connective tissue lineage ^10, 12, 13^. This is consistent that a blastema has a similar gene expression profile to a developing limb bud. Limb bud cells derived from the lateral plate mesoderm can participate in varied connective tissue lineages, but not other non-connective tissues, such as the muscle ^14^ Considering these, dermal fibroblasts may get reprogrammed and become limb bud-like cells. On the other hand, another possibility can explain the emergence of blastema cells from the differentiated dermis. It is well known that multipotent stem cells exist throughout the body. The dermis involves many cells. Thus, it is feasible that the axolotl dermis contains stem cells, and that the stem cells participate in blastema formation. However, molecular and histological descriptions are largely unknown.

The determination of the regeneration induction molecules makes us possible to investigate molecular regulations, which dermis-derived cells receive just after limb amputation. Most blastema cells are derived from the dermis ^10^. And the dermis-derived cells show connective tissues-restricted multipotency in limb regeneration ^10, 12, 13^. The multipotent blastema cells can be induced by B2FF application in axolotls. Hence, investigation of a regulation downstream of B2FF in dermal fibroblasts leads to understandings of cellular reprogramming in amphibian organ regeneration.

Here, we explored the downstream gene network of B2FF by comparing gene expressions *in vitro*. We used mouse dermal fibroblasts as a representative of non-regenerative animals and newt skin fibroblasts as a representative of regenerative animals for the comparative analysis. The following *in vivo* experiments were performed using axolotls because of the benefits of fluorescent observation. The function of *Pde4b* in conferring multipotency was investigated by the usage of a chemical compound and the mosaic Pde4b crispants. Our findings provide important insights into the generation of multipotent cells during limb regeneration in caudate amphibians.

## Results

We first developed an *in vitro* system to investigate the blastemal transformation of dermal fibroblasts. We cannot keep axolotl cells in the satisfyingly proliferative state for a long time. Cultured axolotl cells were inevitable that they decrease in cell division and undergo senescence state after a couple of passages. Dermal fibroblasts derived from newts, however, were highly mitotic and could be stably cultured for a long time. Newt cell line was established from an animal that stably expresses red fluorescent proteins on the cell membrane (Fig. 1A, B, E, F). Due to the property of the membrane-anchored mCherry, the fluorescent signal was observed on the cell membrane and shows the spotted pattern (Fig. 1E, F). The growth rate was approximately 1 doubling per week (Fig. 1G). To investigate the blastemal transformation of the cells, we treated the cultured cells with the regeneration inductive molecules (B2FF: Fig. 1C, D). No significant change was observed after 48 hours by B2FF treatment. However, B2FF treated cells showed large cell aggregates 2 weeks after the treatment (Fig. 1H2, J). No cellular aggregates were observed in the control (Fig. 1H1, I). Sections of the cell aggregates were prepared and gene expression of blastemal marker genes was investigated by *in situ* hybridization. *Msx1* expression was confirmable throughout the cell aggregates (Fig. 1K), and *Msx2* expression was much intense in the surface area (Fig. 1L). The signal of *Prrx1* and *Pea3* were weak but recognizable (Fig. 1M, N) as compared to the control (Fig. 1O). These results suggest that newt skin fibroblasts can react with B2FF under *in vitro* environment.

**Figure 1.**
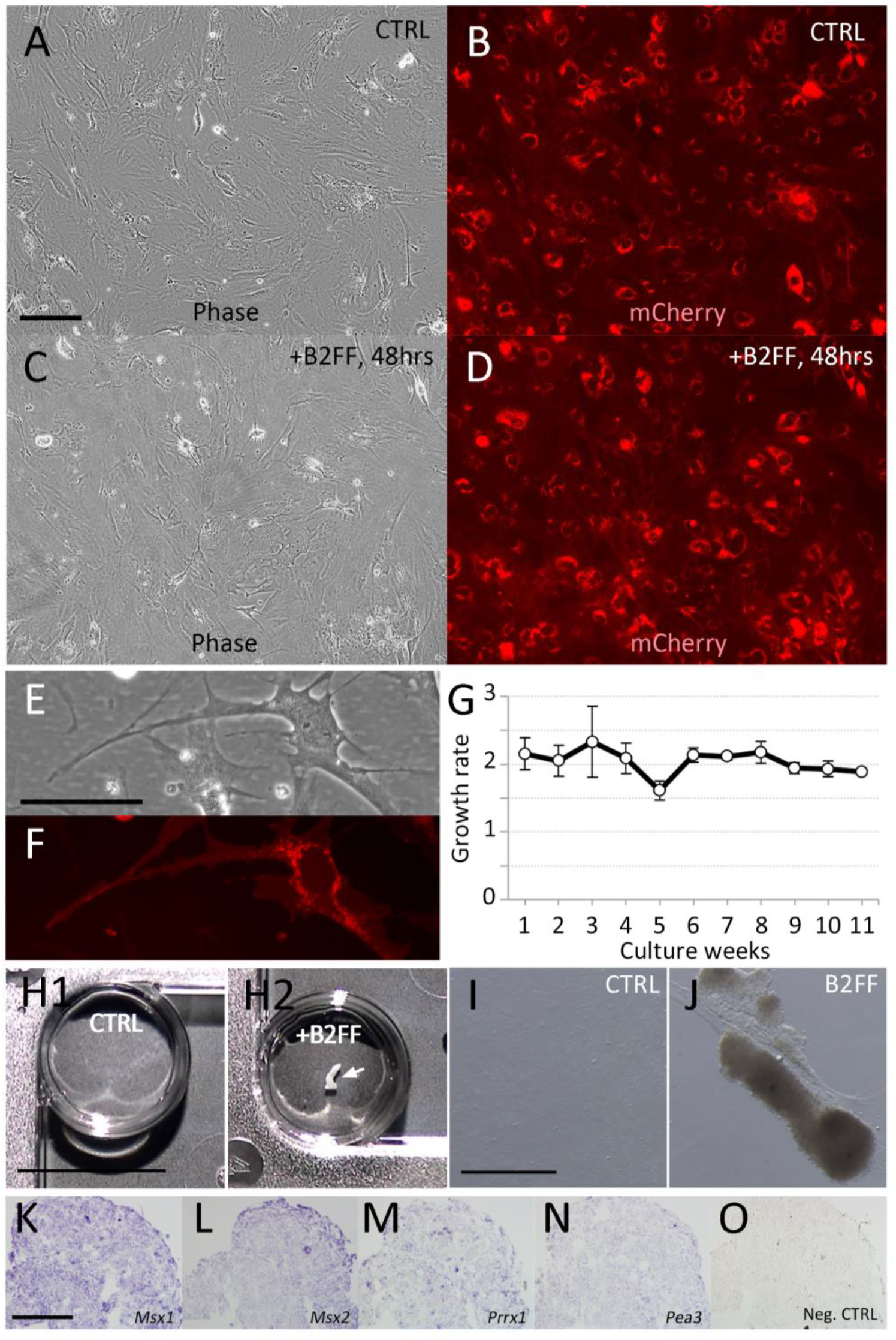
Cultured newt cells and B2FF application. (A, B) Newt cells in the control medium (no B2FF). (C, D) Newt cells 48h hours after the B2FF application. (A, C) Bright-field view. (B, D) Darkfield view. Scale bar in A = 500 μm. (E, F) Higher magnification view of the newt cells in the control condition. Scale bar in E = 100 μm. (G) The growth rate of the cultured newt cells in the control condition. (H1) Newt cells were kept for 2 weeks in the control condition. No cell aggregates could be observed. (H2) Newt cells were kept for 2 weeks in the B2FF-contained condition. Scale bar in H1 = 500 μm. A large cell aggregate could be seen (arrow). (I, J) The bright-field images of the cells in the control (I) and the B2FF-contained (J) condition. Scale bar in I = 200 μm. (K–O) Gene expression pattern of the aggregate formed in the B2FF condition was investigated by *in situ* hybridization. (O) the negative control (no probes). Scale bar in K = 300 μm.

We transgrafted the cell aggregates into an axolotl blastema to investigate the differentiation capability of cells involved in the aggregate. Xenografting of newt cells into axolotl tissues had been described previously ^15^. Newt cells can survive in axolotl tissues for a month. Because of the transparency of an axolotl limb as compared to a newt limb, axolotl limbs are much useful to trace grafted cells in limb regeneration. The B2FF induced aggregate could be easily removed from the plastic dish and transgrafted into an axolotl mid-bud blastema (Fig. 2A). Regarding the control (non-treated), the cultured cells never form aggregates. Hence, cell sheets were collected using a scraper and transgrafted into an axolotl blastema (Fig. 2A). Axolotl limbs are relatively transparent, which allows the trace of the grafted cells by the red-fluorescent (Fig. 2B, C). The grafted newt fibroblasts could survive in the axolotl tissues and participate in a regenerate (Fig. 2B, C). To explore cellular contribution, regenerated limbs were sectioned (Fig. 2D–F). Immunofluorescent analysis revealed the location of the grafted cells (red) and cartilage formation (Col2a; green) in the regenerate (Fig. 2D–F). Little mCherry^+^ was found in the cartilage when the control cells (non-treated) were transgrafted (Fig. 2D1-3). In contrast, the mCherry^+^ cells were identifiable within the cartilage region when the B2FF treated cells were grafted (Fig. 2E1-3). The mCherry signals were always weakened in the cartilage due to the cartilage properties. Cell surface-located and spotted red signals were observable, consisting of the fluorescent pattern observed in Fig. 1F (Fig. 2F). Furthermore, it was suggested that dermis-derived cells can redifferentiate into various connective cell types ^10, 12, 13^. To clarify whether mCherry^+^ cells were observed in other connective tissues, we focused on the tendons. Tenascin is a marker of tendons and ligaments and is expressed in the connecting region between skeletal structures and muscles in amphibians ^16^. Tenascin expression was visualized by immunofluorescence (Sup. Fig.1). Tenascin expression was observed in the peripheral region of the epiphysis and some fibroblast near the epiphysis (Sup. Fig.1A). Confocal observation confirmed that tenascin was located by the mCherry^+^ cells, suggesting that the mCherry^+^ cell became the tendon cell (Sup. Fig.1B, C). Those data suggest that the grafted B2FF treated newt fibroblasts participated in some connective tissue lineage.

**Figure 2.**
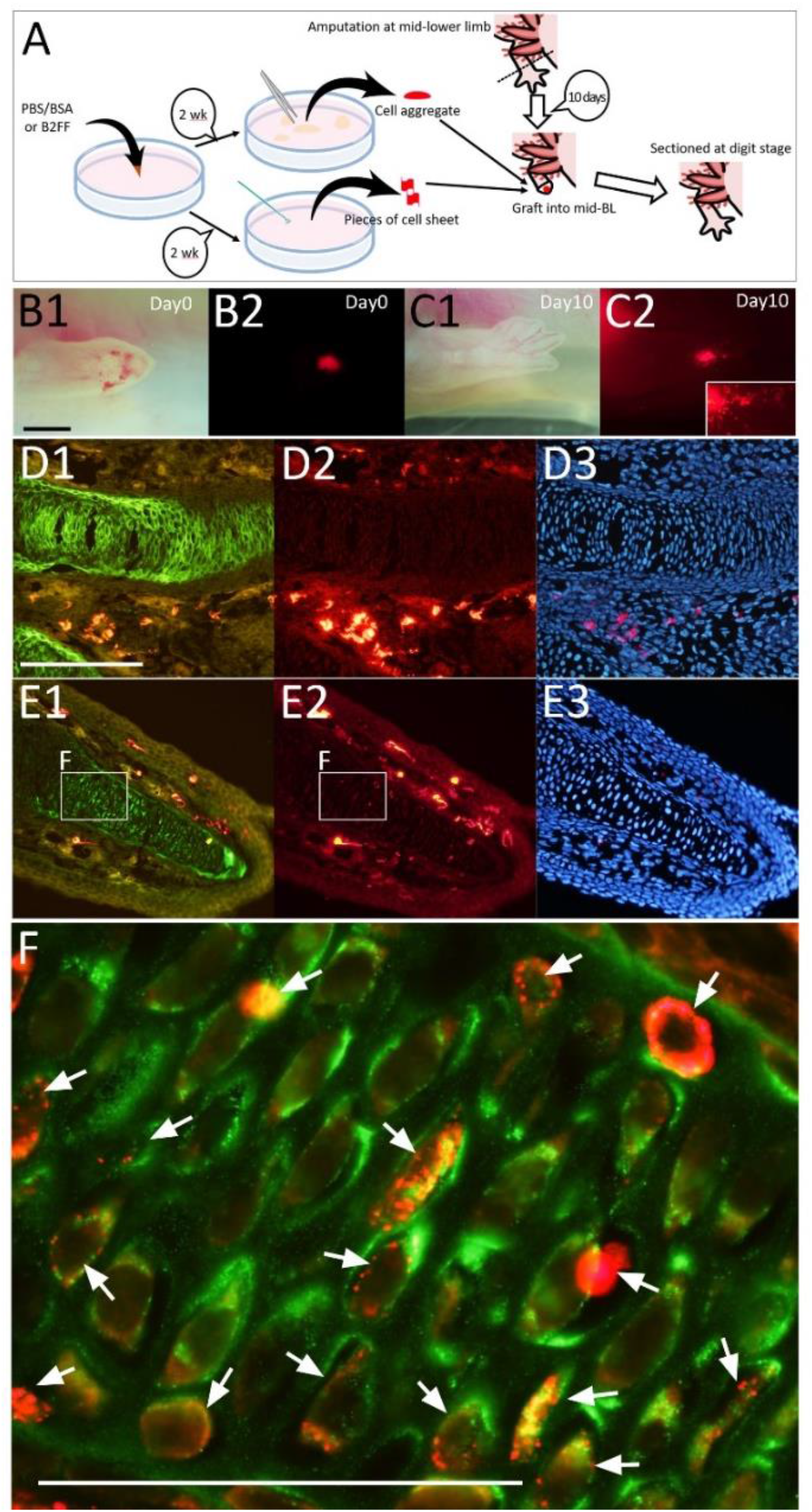
The cultured newt cells were grafted into regenerating axolotl limb blastema. (A) The schematic diagram of the experiment. (B) Just after the grafting. Scale bar in B1 = 1 mm. (C) 10 days after the grafting. Newt cells (red) were spread out in the regenerate (insert). (D–F) The distribution of the newt cells was investigated on the section. Immunofluorescent analysis for Col2a (green; D1 and E1) and mCherry (red; D2 and E2) was performed. Nuclei were visualized by Hoechst33342 (blue; D3 and E3). (F) the merged image of the boxed region in E1 and E2. The red signals were observable within the Col2a+ region. The scales bar in D1 and F are 300 and 100 μm, respectively.

Next, the comparative RNA-Seq analysis was conducted in order to identify the genes related to the acquisition of multipotency. To investigate gene dynamics prior to the formation of cell aggregate, cells 48 hours after B2FF treatment were harvested for comparison. Furthermore, to focus on the specific gene dynamics in the regeneration-competent animals, mouse cells were used as a comparison. Mouse dermal fibroblasts were collected from neonates and 5 passages were undergone before the experiment. B2FF treatment was performed for 48 hours. RNA-seq was performed by CAGE-seq. To compare gene expression between newts and mice, gene symbols were used in this comparison. This was because of the lack of fixed genome information of *Pleurodeles waltl*. Thus, a precise orthologue determination could not be fixed. We found 4185 genes were commonly expressed in the cultured cells of both species (Fig. 3A, Supplemental Data 1). Apparently, no huge different profiles could be found as both species were compared (Fig. 3A). B2FF treatment did not change the gene profile greatly as shown in Fig. 3A. Among those genes, we selected the genes showing FC<-0.5 and FC>0.6. We found that there were 42 genes affected by B2FF treatment (Fig. 3B, Supplemental Data 1). We found out that *Col1A2*, which is a major component of the dermis, was down-regulated by B2FF treatment in both species (Supplemental Data 1). This is consistent with the report, in which an axolotl limb blastema is a less collagenous structure ^17^. We further focused on the genes which react oppositely to B2FF in the two species (Fig. 3B). *Fblim*, *Loxl2*, *Pde4b*, *Spry1*, *Timp3* were upregulated in newt cells and down-regulated in mouse cells by B2FF treatment. *Adra2a* showed opposite dynamics. To confirm the RNA-seq results, quantitative RT-PCR (qPCR) analysis was performed (Fig. 3C). The gene expression profiles were consistent with the results from RNA-seq. (Fig. 3C).

**Figure 3.**
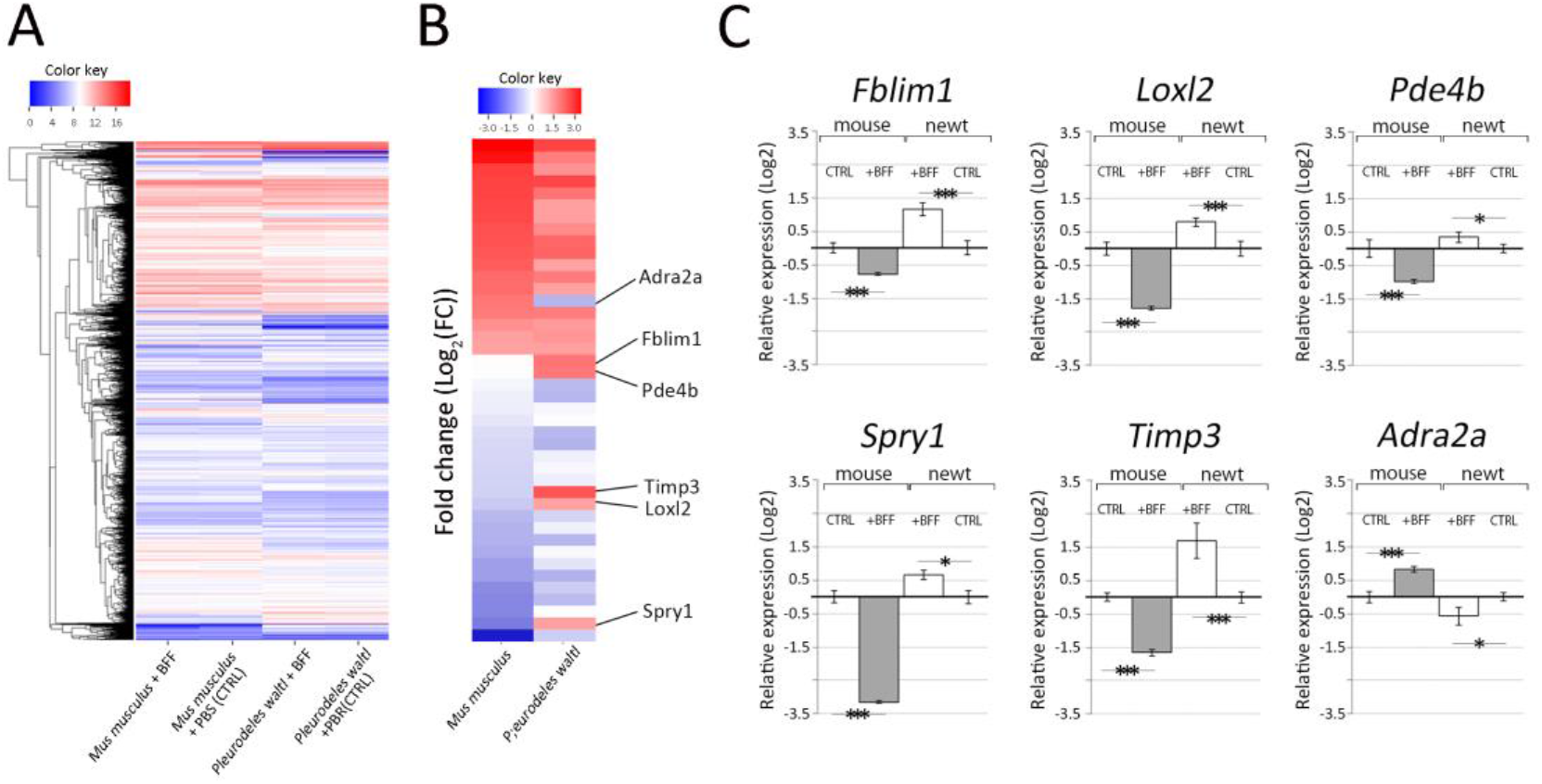
Selection of genes responding to B2FF in cell culture. (A) Z-score obtained via CAGE-seq is visualized as the heatmap. The samples are mouse dermal fibroblasts with B2FF application, mouse dermal fibroblasts without B2FF application (CTRL), newt dermal fibroblasts with B2FF application, and newt dermal fibroblasts without B2FF application (CTRL) from the left. (B) the heatmap of the selected genes, which were up/down-regulated by B2FF application. (C) the gene expression pattern was confirmed by the quantitative RT-PCR. Genes, which were oppositely responded to B2FF in mouse and newt cells, were selected and confirmed. *p<0.05. ***p<0.01.

These 6 genes dynamics were also investigated in axolotl blastemas (Fig. 4). As mentioned, axolotls have many advantages *in vivo* experiments. Thus, we investigated the gene expression patterns in axolotl blastemas in order to perform further experiments in axolotls. qPCR analysis revealed that *Fblim*, *Loxl2*, *Pde4b*, *Spry1*, *Timp3* were upregulated in the blastema. *Fblim1*, *Loxl2*, and *Pde4b* showed quick upregulation after limb amputation (Fig. 4). Only *Pde4* expression was settled down at 10 days post-amputation (dpa). *Spry1* and *Timp3* have a late-activation profile in axolotl limb blastemas. *Adra2a* expression was not detectable throughout the period we tested. These gene expression patterns were confirmed by *in situ* hybridization. (Fig. 5). Consistently, *Adra2a* was not detectable at 2, 5, and 10 dpa (Fig. 5A, A’, G, G’, M, M’). *Fblim1* expression could be recognized in the amputation region from 2 dpa to 10 dpa (Fig. 5B, H, N). The signal of *Fblim1* could be observed at the border of the amputation plane (Fig, 5B’), but the signal was weakened in the later time points in the stump regions (Fig. 5H’, N’). *Loxl2* expression could also be detected from 2 dpa (Fig. 5C, I, O). The *Loxl2* expression was confirmed in the part of the basal layer of the blastema epithelium (Fig. 5C, I). *Loxl2* activation was relatively specific in the distal (blastemal) region and not apparent in the stump region (Fig. 5C’, I’, O’). Upregulation of *Loxl2* likely takes place in the specific cells, rather broader blastema mesenchymal cells. *Pde4b* was upregulated in the early stages (2 and 5 dpa; Fig. 5D, J). The upregulation of *Pde4b* could also be observed around the amputation sites 2 and 5 dpa (Fig. 5D’, J’). But *Pde4b* expression was down-regulated in the later stages (10 dpa) (Fig. 5P, P’). *Spry1* signals were not obvious 2 dpa in both the distal and the proximal region (Fig. 5E, E’). But the signal became confirmable in the blastema 5 and 10 dpa in the blastema (Fig, 5K, Q). *Spry1* expression was also observed around the amputation site 5 dpa (Fig. 5K’). No signal was confirmed in the stump region 10 dpa (Fig. 5Q’) Upregulation of *Timp3* could be detected 2 and 5 dpa (Fig. 5F, L). However, the *Timp3* signal was weakened in the proximal region of the blastema although the distal blastema maintains a relatively high level of *Timp3* expression at 10 dpa (Fig. 5R). The faint signal could also be observed around the amputation site 2 dpa (Fig, 5F’). But no signal could be detected in the proximal region 5 and 10 dpa (Fig. 5L’, R’) Those data suggest that the selected 5 genes from the RNA-seq in mouse and newt cells were consistently upregulated in the axolotl blastema.

**Figure 4.**
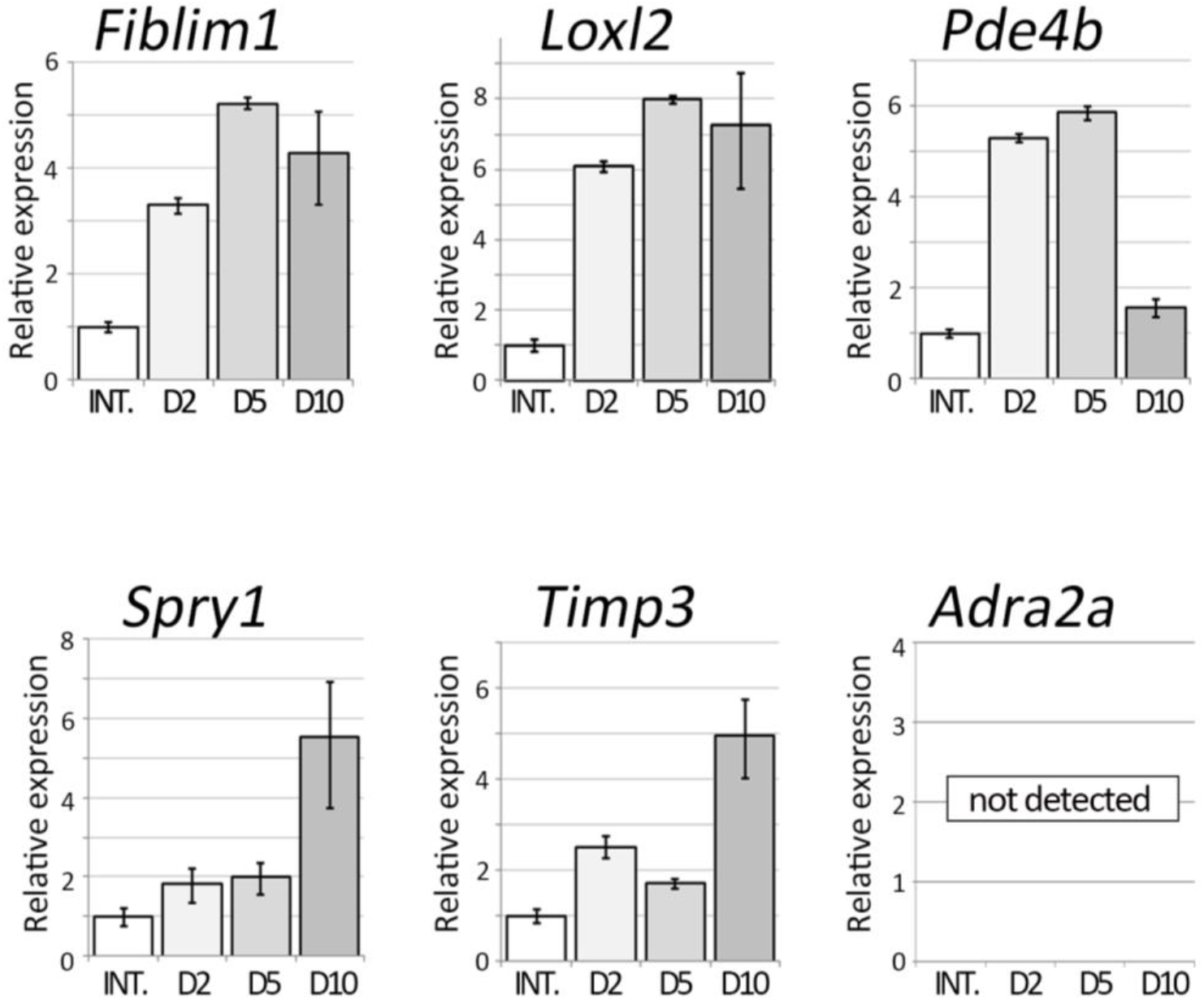
Gene expression patterns of the selected genes. The genes selected from the comparative analysis between the mouse and newt cells were investigated in axolotl blastemas. The quantitative RT-PCR to the samples, which were prepared from blastemas at different time points, was performed. INT. = intact limbs. D2 = 2 days after limb amputation.

**Figure 5.**
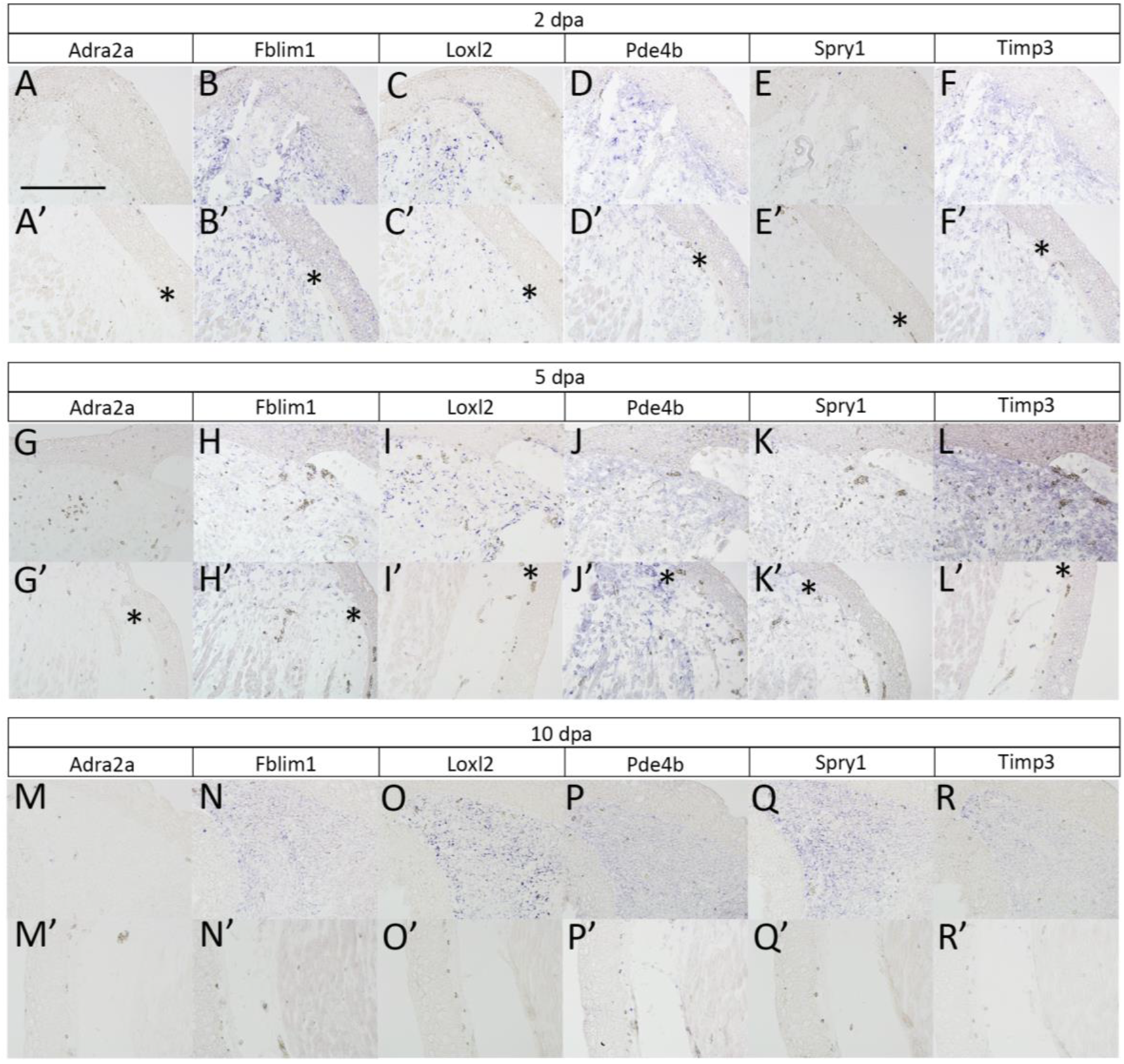
Gene expression patterns revealed by *in situ* hybridization. The blastema samples were prepared at 2 dpa (A–F), 5 dpa (G–L), and 10 dpa (M–R). (A, G, M) expression pattern of *Adra2a*. (B, H, N) expression pattern of *Fblim1*. (C, I, O) expression pattern of *Loxl2*. (D, J, P) expression pattern of *Pde4b*. (E, K, Q) expression pattern of *Spry1*. (F, L, R) expression pattern of *Timp3*. The upper panels (A–R) show the distal region of the amputated limbs. The lower panels (A’–R’) show the proximal or the approximate border of the amputation plane. The asterisks indicate the disconnection of the dermal collagen layer, suggesting the amputation plane. Scale bar in A = 400 μm.

Instability of cellular differentiation should be induced in the early phases prior to blastema formation. Blastemas at 5 dpa have been reported to express blastema marker genes and blastemas at the time point sometimes take a dome shape containing blastema cells ^18^. Moreover, the induction mechanism of cellular dedifferentiation and maintenance mechanism of an undifferentiated state would be different. Considering those, genes related to the induction of instability of cellular differentiation would be upregulated in early, and down-regulated in later. From this point of view, we thought *Pde4b*, which showed early upregulation and late down-regulation, was suitable for this criteria.

We next attempted to inhibit *Pde4b* functions in limb regeneration using a chemical inhibitor, Rolipram. Rolipram binds to the catalytic sites of PDE4B at several amino acids, where are 100% conserved between human PDE4B and axolotl PDE4B (Sup. Fig. 2). It is well known that PDE4B has a function to hydrolyze cAMP to 5’AMP ^19^. To confirm the inhibitory effects to PDE4B in axolotl tissues, cAMP concentration in axolotl limbs was measured by ELISA (Fig. 6A). Consistently, cAMP concentration in limb tissues was upregulated by the 7 day-Rolipram treatment (Fig. 6A). This suggests that Rolipram can effectively inhibit hydrolysis of cAMP by PDE4B in axolotl limbs. Next, we investigated the dermal fibroblast’s transdifferentiation into cartilaginous cells in the presence of Rolipram (Fig. 6B–E). The limb skin from a GFP animal was transgrafted onto a normal animal, and the grafted limb was kept a week for the recovery from the grafting damages (Fig. 6B). Then, the limb was amputated, and the limb-amputated animals were kept in Rolipram-containing water until digits were identifiable (Fig. 6B, C1-C3). The GFP positive domain was expanded from the amputation stump to the digit tips of the regenerate (Fig. 6C1-3). The regenerates were fixed and sectioned. *Col2A1*^+^ cartilage cells were revealed by *in situ* hybridization and the location of the GFP^+^ cells was revealed by immunofluorescence (Fig. 6D, E). In the control samples, GFP^+^ cells were observable in the *Col2A1*^+^ region (Fig. 6D, Table 1). Five limbs were obtained and sectioned. Sectioning was performed on the entire limb along the dorsoventral axis. Only sections with GFP^+^ cells in the mesenchymal region on the prepared sections were extracted, and the GFP^+^ cells were counted. We found 3321 GFP positive cells in 104 extracted sections, of which 149 GFP positive cells were *Col2A1*^+^. In the Rolipram-treated animals, GFP^+^ cells were little observed in the *Col2A1*^+^ region (Fig. 6E, Table 1). We obtained 36 sections from 7 Rolipram-treated samples. Although the number of sections obtained from a single limb sample was about the same as the control, the number of sections to be extracted in the Rolipram-treated samples was lower as compared to the control. This was because of the poor participation of GFP+ cells in the regenerates compared to controls. Using the same method of cell counting as the control, we observed only one *Col2A1*^+^ cell out of 462 GFP^+^ cells (Table 1). It is also noteworthy that cartilage formation in regenerates was not influenced by the Rolipram treatment. These suggest that Rolipram treatment increases cAMP concertation in tissues resulting in suppression of the transdifferentiation from dermal fibroblasts to cartilaginous cells.

**Figure 6.**
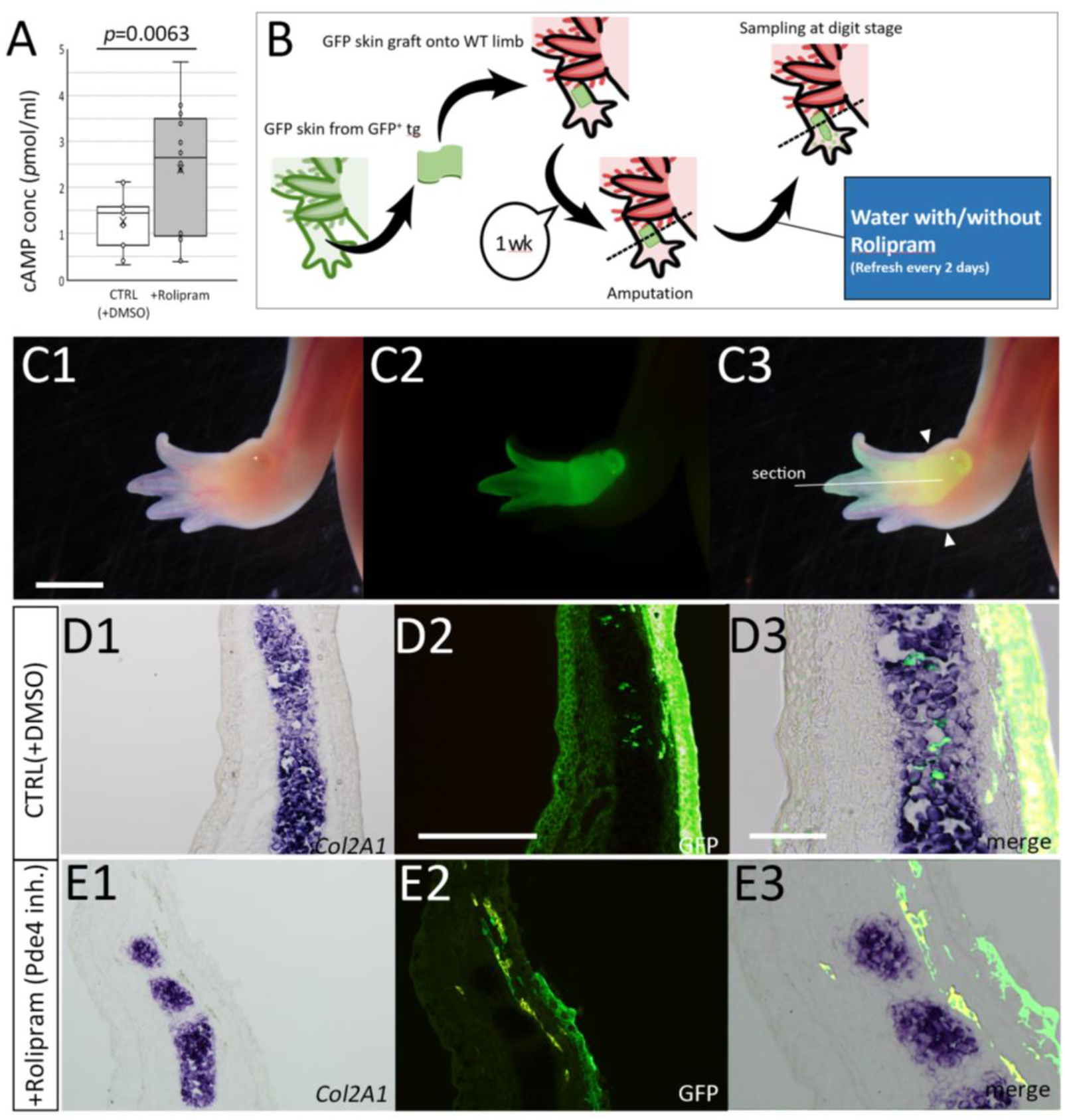
Impaired transformation of dermal fibroblasts into cartilage cells by inhibition of Pde4b function by Rolipram. (A) Cyclic AMP concentration in limb tissues was measured by ELISA. (B) The schematic diagram of the experiment. (C) Rolipram treated limb in the bright-field view (C1), the dark field view (C2), and the merged image (C3). Scale bar in C1 = 3 mm. (D, E) Distribution of the GFP^+^ cells in the regenerated limbs in the control (D) and the Rolipram treated animal (E). Col2a1 expression was visualized I by *in situ* hybridization (D1, E1) and GFP signals were detected by immunofluorescence (D2, E2) on the identical sections. D3 and E3 were the highly magnified and merged images. The scale bar in D2 and D3 are 300 μm and 100 μm, respectively.

**Table 1:** The cell count of GFP^+^ Col2a1^+^ cells in Rolipram treated limbs

|  | Number of specimens | Number of pictures | Count for GFP+ Cells | Count for GFP+/Col2A1+ | Ratio(%) |
| --- | --- | --- | --- | --- | --- |
| CTRL(DMSO) | 5 | 104 | 3321 | 149 | 4.49 |
| Rolipram | 7 | 36 | 462 | 1 | 0.22 |

We further investigated increasing in cAMP concentration impaired cartilage transdifferentiation from dermal fibroblasts. Dibutyryl-cAMP is a cell-permeable cAMP analog that activates cAMP-dependent protein kinases ^20^. Similarly, GFP^+^ skin was transgrafted onto a normal animal and the GFP skin-grafted limb was amputated to trace the lineage of GFP^+^ cells in the absence or presence of dibutyryl-cAMP (Fig. 7A). The amputated limbs were kept until the regenerates reached the digit stage (Fig. 7B, F). To visualize GFP and Col2A1, we performed immunofluorescence on the identical sections (Fig. 7C–I). In the control sample, GFP^+^ cells in the cartilaginous region could be detected as well as epidermis and dermis (Fig. 7C–E). On the other hand, the Dibutyryl-cAMP treated limbs showed a little number of GFP^+^ cells in the cartilaginous region (Fig. 7G–I). We plotted the rate of the GFP^+^/Col2A1^+^ in the regenerates (Fig. 7J). We counted 13 sections from 8 independent animals in the control and 21 sections from 17 independent animals in the Dibutyryl cAMP-treated animals. It is noteworthy that the two exceptional plots in the Dibutyryl cAMP-treated samples were derived from an identical animal. These results consistently suggest that cAMP concentration influences a fluctuation of differentiation of dermal fibroblasts.

**Figure 7.**
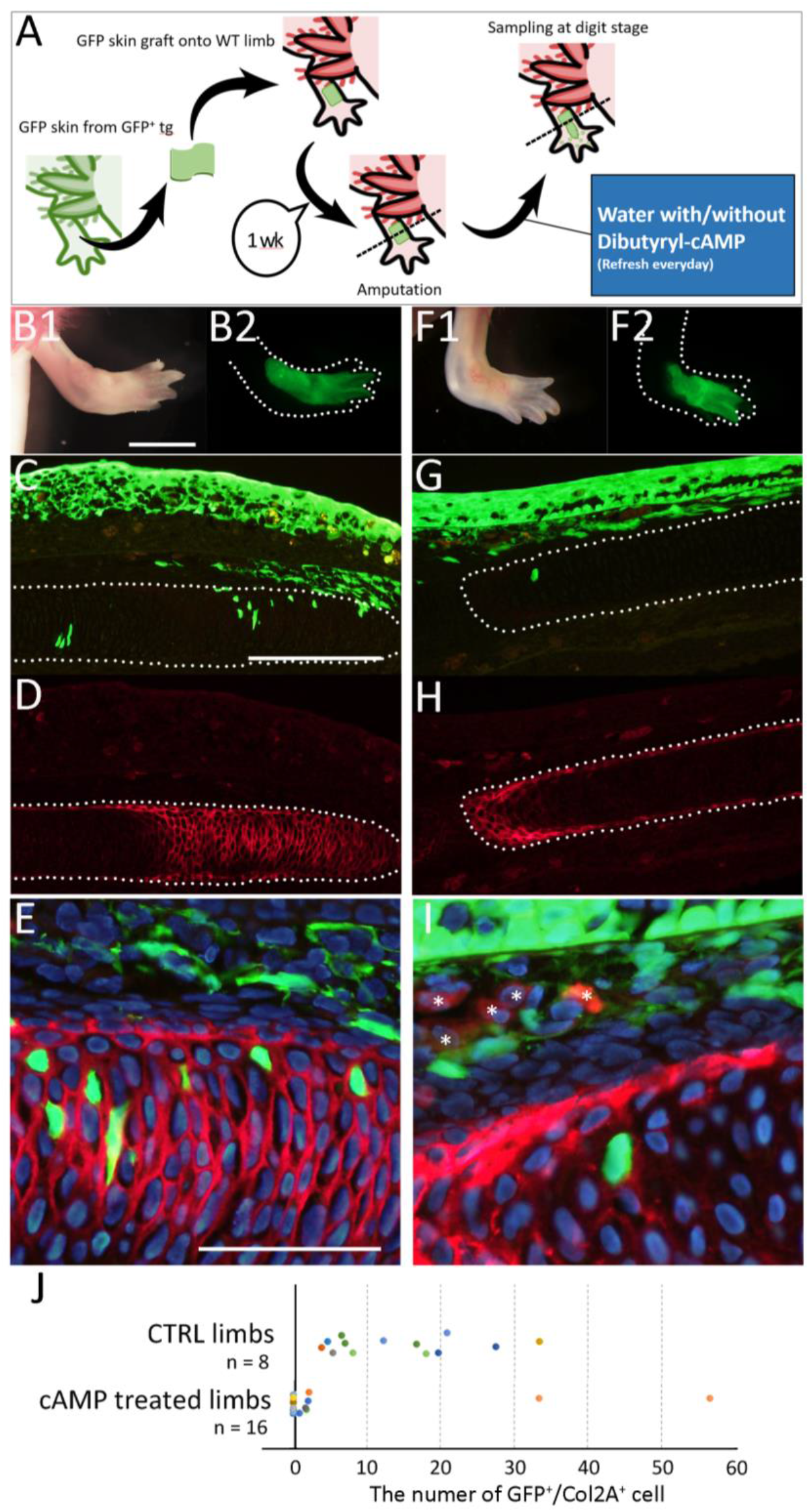
Distribution of GFP+ cells in the dibutyryl-cAMP treated animal. (A) the schematic diagram of the experiment. The GFP-skin grafted limb of the control animal (B–E) and the dibutyryl-cAMP treated animal (F–I). (B, F) The GFP-grafted limb just before sampling. (C–E, G–I) The distribution of the GFP^+^ cells and Col2a expression were visualized by immunofluorescence. (E, I) the merged and higher magnified image of C and D. The asterisks in I indicate the red blood cells. The dotted lines indicate the border of the cartilage. (J) The plot of the number of GFP^+^Col2a^+^ cells. The scale bars in B1, C, and E are 3 mm, 300 μm, and 100 μm, respectively.

Next, we attempted to inhibit PDE4B in cultured newt cells by Rolipram (Fig.8). The newt cells were cultured as above. The control (no B2FF) and the B2FF treated cells gave rise to the sheet and the aggregate formation, respectively (Fig. 8B–D). Rolipram application into B2FF culture media resulted in no aggregate formation (Fig. 8E; n=6/6). We grafted the ^+^Rolipram/^+^B2FF cell as a sheet since no aggregate formation could be obtained (Fig. 8A). The participation of the newt cells into cartilage was assessed at the digit stage. The grafted cells could survive and expand in the regenerate (Fig. 8F, G). The section revealed that a large number of mCherry^+^ newt cells could be observed outside of the cartilages (Fig. 8H, I). Even though the mCherry^+^ newt cells were located just by the regenerated cartilage, no participation of the grafted newt cells in the cartilage could be observed (Fig. 8J). All cell counts were shown in Table 2 and Fig. 8K. These results strongly suggest that Rolipram treatment inhibits re-differentiation from dermal fibroblasts to cartilaginous cells.

**Figure 8.**
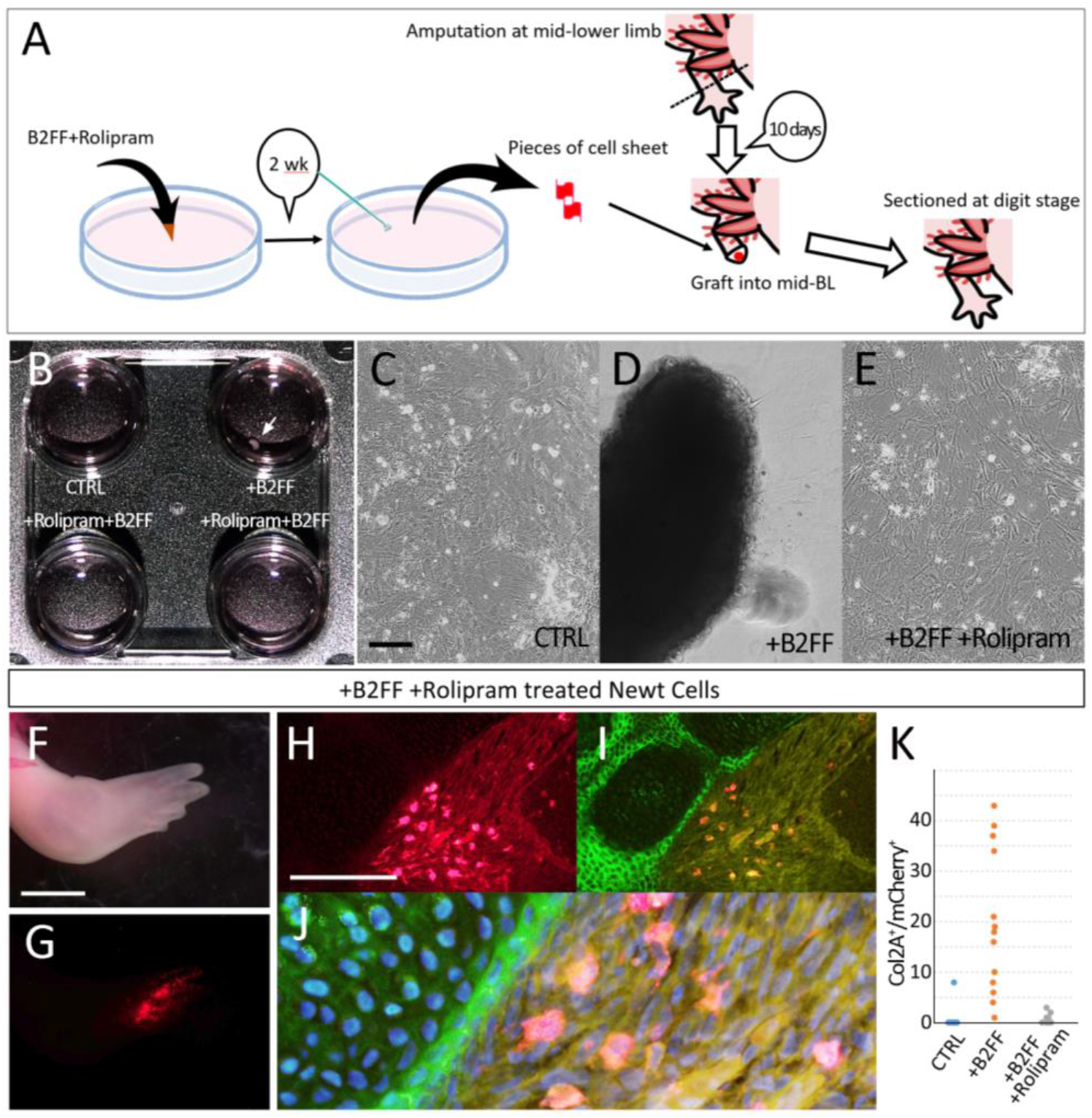
Inhibition of the participation of dermis-derived newt cells into cartilage by Rolipram treatment. (A) the schematic diagram. (B) Rolipram treatment inhibited the formation of cell aggregate by B2FF application. (C–E) Cell morphology 2 weeks after the treatment. (C) the control. (D) The B2Ff treated cell. The aggregate was formed. (E) The Rolipram+B2FF treated cells. No cell aggregate could be seen. (F–J) Distribution of the grafted newt cells in axolotl limb regeneration. (F) The newt cell-grafted limb regenerates normally. (G) The grafted newt cells (red). (H–J) The distribution of the grafted newt cells and cartilages was visualized by immunofluorescence. (J) the high magnified and merged image of the H and I. (K) the plot of the number of the Col2A^+^mCherry^+^ cells. The scale bars in C, F, and H are 500 μm, 2 mm, and 300 μm, respectively.

**Table 2:** The cell count of Col2a1^+^ mCherry^+^ cells in newt-cell grafted axolotl limbs

|  | Number of specimens | number of pictures | Count for mCherry+ cels | Count for Col2A+/mCherry+ cells | Ratio (%) |
| --- | --- | --- | --- | --- | --- |
| CTRL | 7 | 19 | 525 | 8 | 1.5 |
| +B2FF | 4 | 13 | 799 | 256 | 32.0 |
| +B2FF/+Rolipram | 4 | 10 | 331 | 7 | 2.1 |

We next directly manipulated the *Pde4b* gene in axolotls. CRISPR/Cas9 systems allowed to generate mosaic *Pde4b* knockout animals (*Pde4b* crispants). We had not succeeded in generating homogenous *Pde4b* crispants. Five crispants were used and the knockout rate was assumed by ICE-analysis (30-58%, Fig. 9G). We labeled the dermal fibroblasts by GFP electroporation and traced the lineage during limb regeneration (Fig. 9A–E). The electroporation was performed prior to limb amputation. The electroporated limb was amputated 3 days after the electroporation, and the animals were kept until digits were apparent (Fig. 9A, D). In both the control limbs (n=4) and the Pde4b crispant limbs (n=8), GFP^+^ cells could be seen in the regenerated limbs (Fig. 9A, B, D, E). The section revealed that GFP^+^ cells could be seen in the *Col2A1*^+^ cartilage region and other connective tissues in the regenerated autopodial region in both the control animals and crispants (Fig. 9C, F). We counted GFP^+^ cells in the regenerated autopodial region (Fig. 9G). Longitudinal sections were made throughout the regenerate. The GFP^+^ and the GFP^+^*Col2A1*^+^ cells were counted on all sections. The control limbs, in which the knockout score was 0, showed that many GFP^+^ cells differentiated into *Col*2a1^+^ cartilaginous cells. On the other hand, *Pde4b* crispants showed that much fewer GFP^+^ cells participated into *Col2A1*^+^ cartilage. The alignment of the GFP^+^*Col2a1*^+^/GFP^+^ ratio with the knockout score calculated from ICE analysis revealed a strong correlation (R^2^=0.912). Limbs from an animal having a higher knockout score showed a low integration rate of GFP^+^ into a cartilaginous region. In contrast, limbs with a lower knockout score showed a relatively higher integration rate. This suggests that the *Pde4b* function relates to the conversion from dermal fibroblasts to cartilage cells in axolotl limb regeneration.

**Figure 9.**
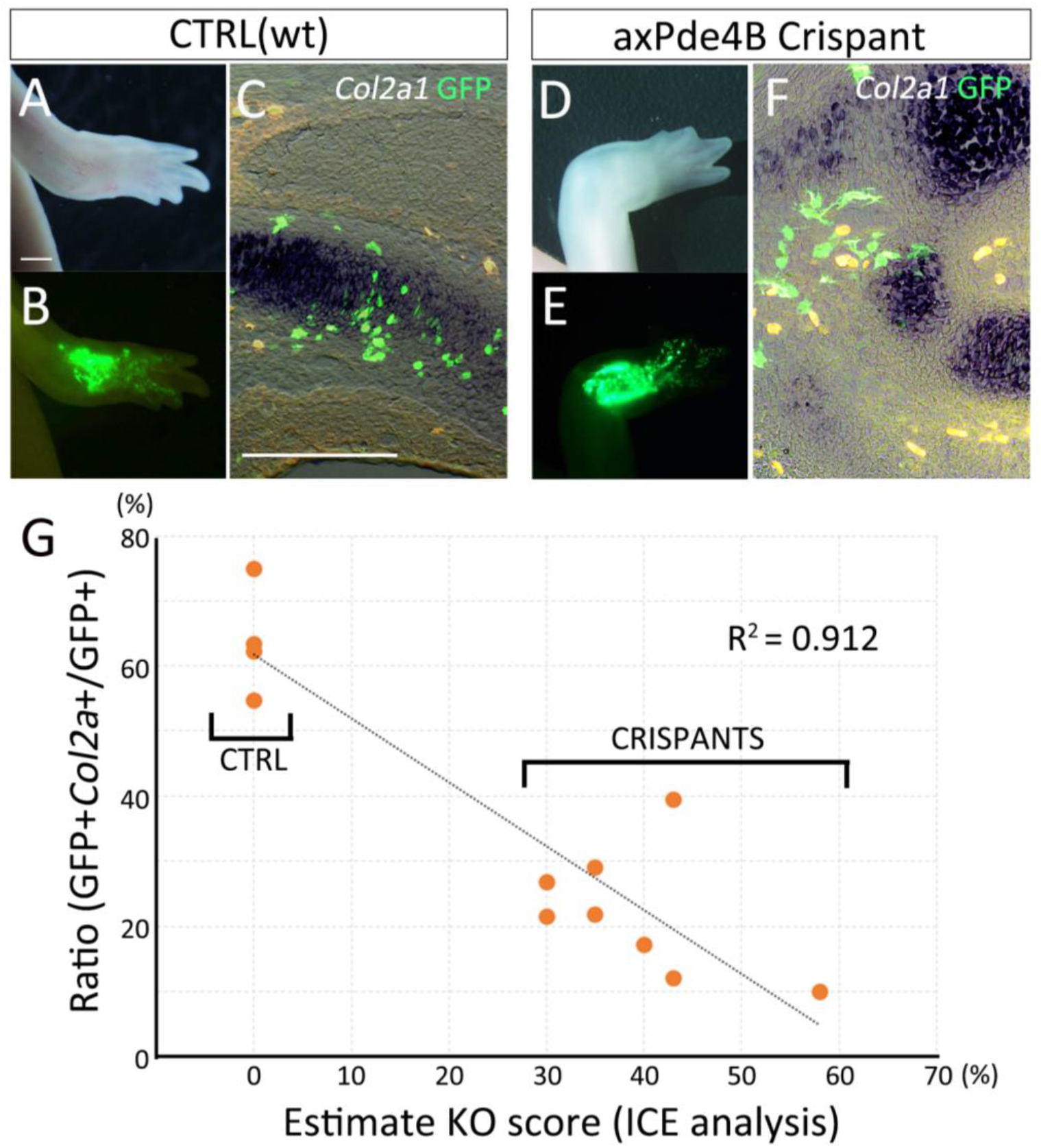
Transdifferentiation from the dermis to cartilage in the mosaic Pde4b crispants. (A–C) the control animals (wt: wild-type). (D–E) The mosaic Pde4b crispants. (A, D) limb morphology. (B, E) The distribution of the GFP^+^ cells. GFP plasmids were electroporated in the dermal region prior to amputation. (C, F) the merged images of the *Col2a1*, GFP, and the bright field view. *Col2a1* and GFP were visualized by *in situ* hybridization and immunofluorescence, respectively. The scale bars in A and C are 1 mm and 300 μm, respectively. (G) the plot of the GFP+*Col2a1*+/GFP+ ratio along with the KO score calculated by ICE analysis.

## Discussions

### The cultured newt cells derived from the dermis

We cultured fibroblasts from a newts’ limb skin (dermis), in which many types of cells exist. It is reasonably assumed that dermal fibroblasts are not homogenous, rather heterogeneous. Moreover, the determination of fibroblasts is still ambiguous. Thus, it is still difficult to determine the cultured cells we used precisely. In the present study, we obtained the constantly dividing fibroblasts (Fig. 1). During the process of establishing the cell line, it is very likely that certain cell populations were selected and survived. The RNA-Seq data revealed that the cultured fibroblasts express *Colla2*, *Vimentin*, and *Twist1* (Supplemental data 1), which are well-known marker genes as fibroblast marker genes. The expression profile reasonably suggests dermal fibroblasts were dominantly cultured in our experiment. However, culturing cells are dividing. Differentiated dermal fibroblasts *in vivo* are assumedly not actively dividing. Thus, our procedures in cell preparation might somehow transform cells. On the other hand, the cultured newt cells derived from the dermis did not show cartilage differentiation when the cells were grafted in the axolotl blastema (Fig. 2). This suggests that the cultured cells were not multipotent and that the culture condition did not provide multipotency. Further characterization of the cultured cells should be necessary to determine for precisely describing the cells we used.

### The xenografting between axolotls and newts

The grafted newts cell could survive and differentiated into a couple of cell types in the axolotl limb (Fig. 2). Such xenografting between an axolotl and a newt could be found in the histology of the amphibian regeneration study ^21, 22^. The grafted tissues and cells functioned physiologically in the xenografted environments. However, it is still unknown that cells from one species precisely behave normally in the other. On the other hand, limb regeneration can be induced by B2FF in both species ^8^. B2FF, which we used in order to induce regeneration responses in urodele amphibians, are the recombinant proteins, whose amino acid sequences are derived from a mouse or a human. Of course, the axolotl B2FF genes can induce limb regeneration reactions when axolotl B2FF are electroporated ^23^. Thus, mouse B2FF has been considered to activate the same or quite similar gene cascades in both species. Even though the initial activation mechanism is identical, it is not known that the following mechanisms are identical or similar. For instance, the time to progress the regeneration stages is different, implying that the grafted newt cells receive inputs from outside at different timing in an axolotl blastema. We are not sure how the differences influence the grafted newt cells and their differentiation.

The reason why we had to use xenografting in the present study is that we cannot find any good way to culture axolotl cells for a long time. There are ways to culture the axolotl cells ^24, 25^. However, it is still tough to have cells that can keep a proliferative state for a long time. To investigate a more focused and detailed mechanism of dermal fibroblasts’ dedifferentiation, finding a way to culture axolotl cells for a long time is needed.

### Gene selection

In this paper, we used a unique method for gene selection. We compared mouse fibroblasts with newt fibroblasts using cultured cells. The comparison would be controversial because mouse cells and newt cells are physiologically different. While acknowledging the differences in various physiological properties, we were able to find out candidate genes by comparing the downstream factors regulated by a common factor, B2FF. In this gene selection, we focused on genes that are inversely regulated by B2FF treatment in mouse and newt cells. This is based on the finding that the dynamics of dermal fibroblasts in the axolotl (limb) and mouse (fingertip) regeneration are different ^26^. Urodeles can induce multipotent cells from the dermis, while mice cannot induce multipotent cells from the dermis. Therefore, we thought that there might be differences in gene expression during the generation of multipotent cells in the early stages of regeneration. However, it is possible that other systems, such as epigenetic regulation, are involved in the process of generating pluripotent cells from the dermis, and further research is needed to determine whether this is reflected in simple differences in transcriptome expression. Although a multidimensional study is definitely necessary, the fact that we were able to find a functional molecule in the comparative analysis between the two species in this study provides a certain amount of positive endorsement for the gene selection method used in this study.

### Pde4b function in transformation of dermal fibroblasts

We found the *Pde4b* gene from our gene selection. *Pde4b* was upregulated in the very early phase in axolotl limb regeneration (Fig. 4, Fig. 5D, J). *Pde4b* was downregulated after blastema cell emergence (Fig. 5P). This expression pattern is reasonable that *Pde4b* has a function in the dedifferentiation stage. Generally, limb regeneration and development share the same or similar gene cascades to form a patterned limb after blastema formation. Considering this, the cellular dedifferentiation process is unique and takes place before blastema formation. Moreover, the functions of cellular dedifferentiation should be down-regulated after blastema formation because blastema cells are going to be re-differentiated. Thus, the *Pde4b* gene expression pattern would be suitable as a factor involved in cellular dedifferentiation.

*Pde4b* encodes an enzyme to hydrolyze cAMP to 5’AMP. Thus, PDE4B functions in cAMP regulation in limb regeneration. It is well known that there are cAMP-dependent pathways, such as the PKA-pathway ^27^. Thus, disturbing PDE4B leads to influencing many intracellular signaling cascades.

Downregulation of cAMP in the very early phase of limb regeneration is likely important to fluctuate cellular differentiation in dermal fibroblasts. This is consistent with the previous report, in which a low level of cAMP in limb blastemas within 7 days after amputation was described ^28, 29^. Functions of the low level of cAMP at the beginning of limb regeneration have not been investigated. Our results suggest that a low level of cAMP contributes to fluctuating cellular differentiation in limb regeneration. On the other hand, cAMP-dependent activities are unlikely to have severe influences on limb regeneration. PDE4B inhibition by Rolipram resulted in increasing in cAMP concentration in tissues and impairing the rate of transdifferentiation from dermal fibroblasts to cartilaginous cells. However, no skeletal pattern defects were observed. Previous reports strongly suggest that organ-level regeneration can be achieved by lineage-restricted cells, with no transdifferentiation ^13^. Considering this, it is reasonable that a fluctuation of cellular differentiation has little influence on the outcome of limb regeneration. Although the fluctuation of cellular differentiation does have little influence on limb regeneration, the mechanism of induction of multipotent cells from differentiated tissues has been valuable. Blastema cells raised from dermal fibroblasts have been considered to be similar to limb bud cells. The connective tissue lineage-restricted multipotency is just like that of lateral place mesoderm derived limb bud cells. Thus, a mechanism of acquisition of multipotency in dermal fibroblasts might be closely related to the reactivation of developmental programs. We believe that reprogramming of cellular differentiation and/or rewinding developmental programs can be learned from amphibian organ regeneration.

## Acknowledgements

We are grateful Ms. R. Hori for taking a role in cell counting and Ms T. Satoh for general support for conducting the experiments. This work was supported by AMED (18bm0704006h0003) and JSPS Grant-in-Aid for Scientific Research (B) (20H03264).

## Materials and methods

### Animals and Cell culture

Axolotls (*Ambystoma mexicanum*) with a nose-to-tail length of 4–8 cm were used. Small axolotls (3–4 cm) were used for chemical treatment. The GFP axolotls were from the Ambystoma Genetic Stock Center (AGSC, Univ. of Kentucky). The axolotls were housed in aerated water at 22 °C. Photoshop CS5 software (Adobe, San Jose, CA, US). Newts (Pleurodeles waltl) were obtained from Hiroshima University (Amphibian research center)^30^. All animal experiments were conducted following the guidelines of Okayama University.

The cells from a TgSceI (CAG::TVA-mCherry-p2A-Cas9) were isolated to obtain mCherry positive cells. Mouse cells were from the skin of C3H newborn mice (1 day old). Skin fibroblasts were isolated as follows. The limb skin was removed from a mCherry^+^ animal by forceps and micro scissors. The skin was minced well and incubated in 0.5% collagenase solution (0.5% collagenase (Wako-Fuji film, Osaka, Japan) in 70% phosphate-buffered saline (PBS)) for 6 hours. Then, the equal amount of trypsin solution (0.5% Trypsin (Wako-Fuji film, Osaka, Japan) in PBS) was added and incubated for 30 minutes. The solution was filtered by 45 μm mesh and enzymatic activity were neutralized by adding the culture medium (10% FBS, 40% Glutamax DMEM (ThermoFisher, MA, USA), 50% water, 300 μg/ml Gentamycin (Nakarai tesque, Kyoto, Japan), and 10 mM HEPES, pH 7.5). A couple of centrifugation was taken for wash, and then the cells were cultured on the plate. FGF2 (#3139), FGF8 (#423-F8), and BMP2 (#355-BM) (R&D Systems, MN, USA) were used and added to the medium at 0.1ug/ml concentration. Cell count for calculating the growth rate was performed by Countess II (Thermo Fisher).

### CAGE-seq and RT-PCR

CAGE-seq was performed by DNAFORM (Kanagawa, Japan). Total RNA was extracted from the cells, which were treated with PBS or B2FF for 48 hours. The samples were sent to DNAFORM (Kanagawa, Japan) in order to perform CAGE-seq. High-quality total RNA was prepared by Trizol (Invitorgen). Mapping of CAGE-tag sequences to the transcriptome assembly of Pleurodeles waltl (Trinity_Pwal_v2.fasta.gz, from iNewt website) was performed by Bowtie2. The read count was through the program featureCounts. The differential analysis was using DESeq2.Heatmap was described by Python. The quantitative RT-PCR was performed using primers listed in the supplemental Table 1. The RNA samples were prepared from 2 independent samples. The quantitative RT-PCR analysis was performed by StepOne^™^ (TermoFisher) and KAPA DNA polymerase (#KK4600, NIPPON genetics, Tokyo, Japan).

### Pde4b knockout by CRISPR/Cas9 and ICE analysis

The animals, in which the *Pde4b* gene was heterogeneously edited, were generated as described ^30^. The two guide RNAs were prepared as follows; gPde4b ver1: GGAGGAGCTGGACTGGTGCC, gPde4b ver2: GTCCGTGTGCTTGTTTGCAG. The guide RNA and sgRNA were purified using a CUGA gRNA synthesis kit. The synthesized gRNA was incubated with Cas9 proteins (IDT) for 30 minutes at room temperature before injection.

Gene knockout score was calculated by ICE analysis (https://ice.synthego.com/#/). The axolotl genome for the ICE analysis was extracted from the limb skin. The small piece of the forelimb skin was removed from an upper arm and the genome was purified using the QIAGEN genome extraction kit (#69506). The standard Sanger sequence was performed to the amplified genomic PCR fragment using the following primers; for: ATGATGAAGGAGCACTGCCCCACC; rev: CTTGTTCGATGCCATCTCGCTGACG.

### Electroporation

Electroporation was performed as previously described ^9^. Briefly, pCS2-AcGFP plasmids (1 μg/μl)) were injected underneath of the skin. The electrodes were placed as nipping the injection site. The electroporation was performed under the following condition; 20V, 50ms pulse, 950 ms interval, and 20 times). The electroporated animals were kept for 3 days and checked the fluorescent signals were under a microscope.

### Cell Grafting

Cultured newt cells were grafted into axolotl limbs. Cell aggregates or sheets were from 14-days cultured wells. Aggregated were formed in the B2FF medium. The aggregates were easily removed from the plastic dish by forceps. Regarding to the wells filled by the regular medium, no aggregates were formed. Cell sheets were isolated by a cell scraper. The isolated cell aggregates or sheets were transferred into an axolotl blastema that was grown for 10 days. The grafted limbs were raised until digits were formed.

### Inhibitor treatment

Rolipram (Tokyo Chemical Industry, Tokyo, Japan) was dissolved in dimethyl sulfoxide (DMSO; Nacalai Tesque, Kyoto, Japan), to prepare a 130 mM stock solution. For inhibitor treatment, we kept the animals in the presence of Rolipram (130 μM) or DMSO, in water. The water was changed every day until the samples were fixed.

### Sectioning, histological staining, and in situ hybridization

Tissue samples were fixed with 4% paraformaldehyde/PBS for 1 day at room temperature, and the fixed tissues were treated in 30% sucrose/PBS at 4 °C for 1 day. Then, the samples were embedded in the O.C.T compound (Sakura Finetech, Tokyo, Japan). Frozen sections of 14-μm thickness were prepared using a Leica CM1850 cryostat (Leica Microsystems, Wetzlar, Germany). The sections were dried under an air dryer. For histological observation, standard trichrome staining was performed using the Trichrome Stain (Masson) Kit (Sigma-Aldrich, St. Louis, MO, USA). The stained sections were mounted using Softmount (Wako Pure Chemical Industries, Osaka, Japan). Immunofluorescence on the sections was carried out according to previously reported methods (Satoh et al., 2007). Anti-GFP (#594, MBL, Tokyo, Japan, 1:500), anti-Col2 antibody (II-II6B3, DSHB, IW, USA, 1:200), anti-tenascin C antibody (MT1, DSHB), anti-rabbit IgG Alexa 488 (ab150077, 1:500) and anti-mouse IgG Alexa 594 (A21203, 1:500; Invitrogen, CA, USA) were used for the immunofluorescence procedure. For the Col2 immunofluorescence, antigen retrieval was necessary (Proteinase K (5μg/ml) for 30 minutes at room temperature). Images were captured using an Olympus BX51 system (Olympus Life Science, Tokyo, Japan). Nuclei were visualized by Hoechst 33342 (Wako-Fuji film, #346-07951). RNA probes and *in situ* hybridization procedures have previously been described (Makanae et al., 2014). Cell counts on the images were performed by Photoshop CS6 software (Adobe).

### ELISA

ELISA was performed using Cayman cyclic AMP ELISA kit (#581001, Cayman Chemical, MI, USA). Samples were prepared from the Rolipram treated animals and control (DMSO treated) animals. Limbs were isolated from the animals and used for the analysis. We set the wavelength at 410 nm, and the absorbance was measured by the microplate reader (CORONA, MTP-880lab). Six limbs of each treatment were independently measured.

## Supplemental Figure legends

**Supplemental Figure 1.**
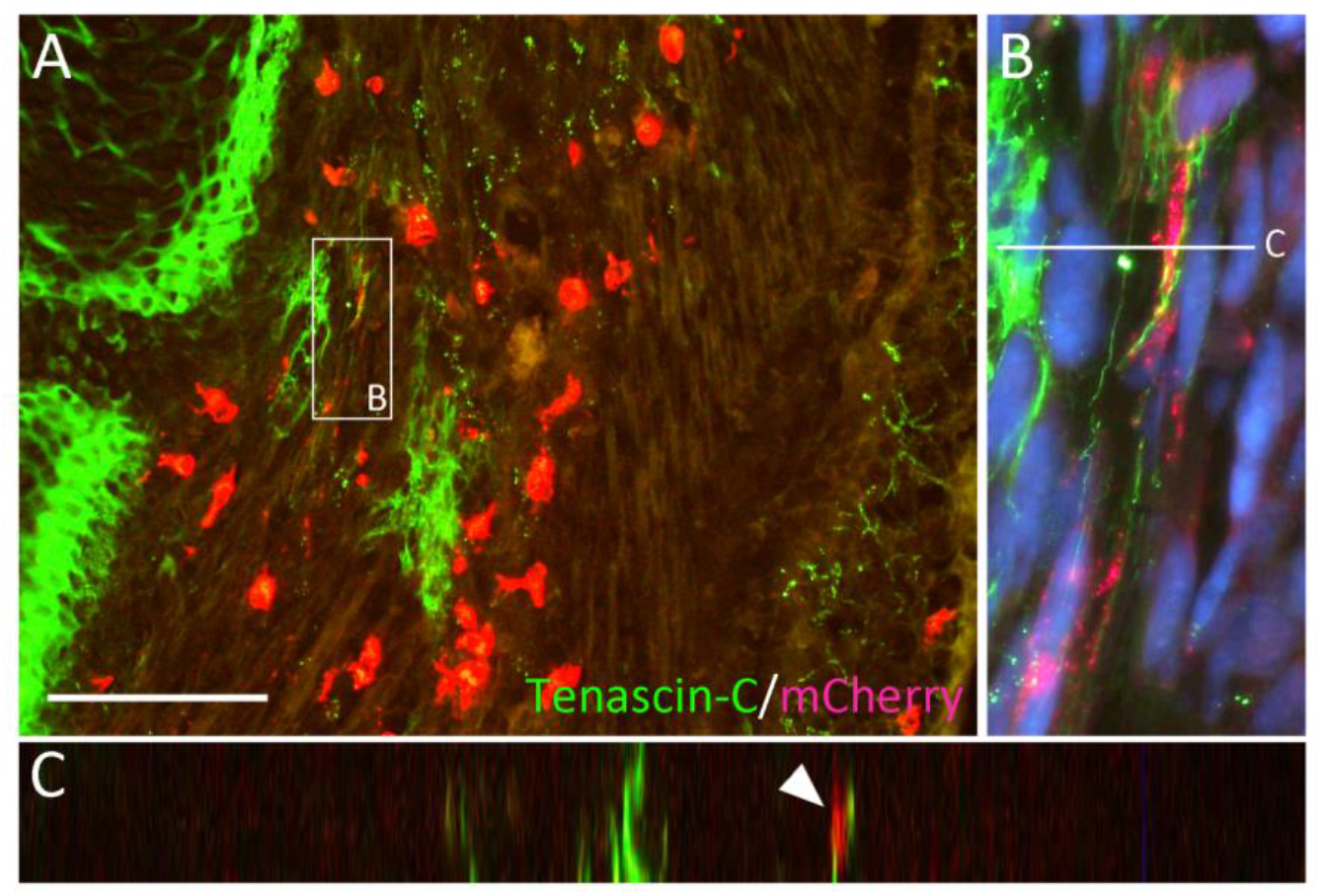
The distribution of the mCherry+ newt cells in the axolotl regenerated limb and Tenascin-C expression. (A) The signals of mCherry and Tenascin-C were visualized by immunofluorescence. The scale bar in A is 200 μm. (B) The higher magnification view of the boxed region in A. The blue color indicates the nuclei. (C) The side view of the region is indicated by the line in B. The Tenascin-C signal is located at the side of the mCherry+ cytoplasm (arrowhead).

**Supplemental Figure 2.**
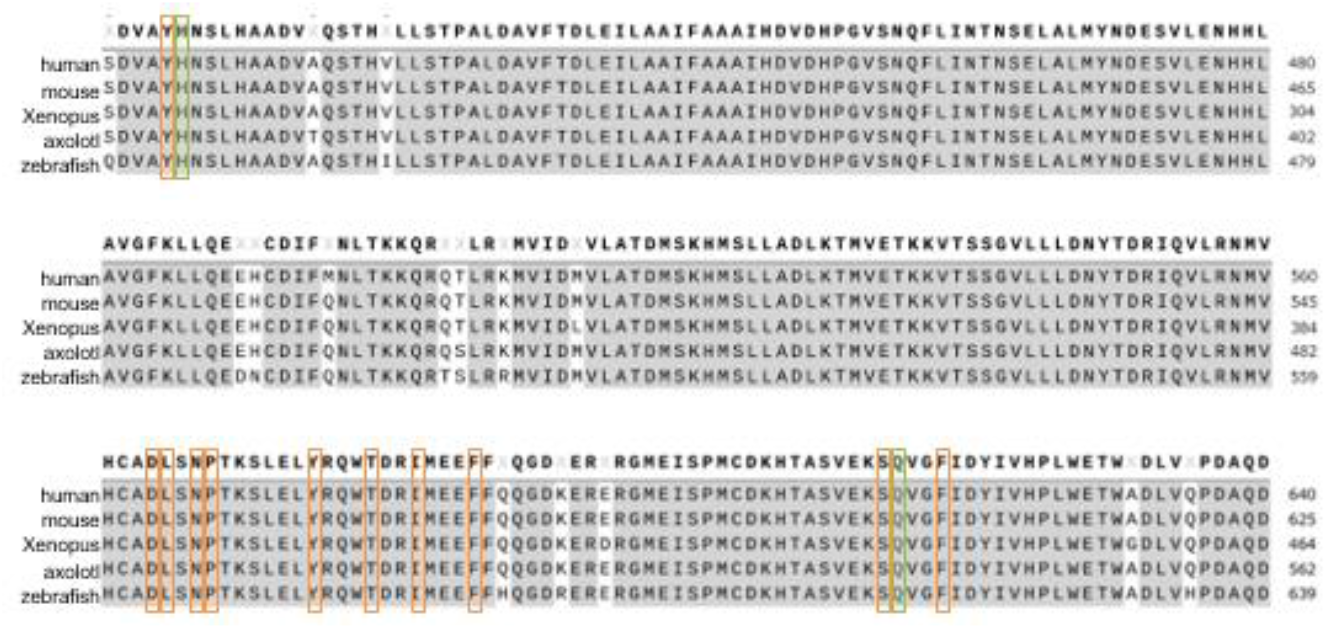
The comparison of the amino acid sequences of PDE4B. The amino acid sequence of PDE4B shown in the figure indicates the nearby region where Rolipram binds. The amino acids surrounded by green lines indicate the residue to which Rolipram forms a hydrogen bond. The amino acid sequence boxed with the orange line indicates the non-ligand residue involved in hydrophobic contact(s).

